# Evaluating large language models in biomedical data science challenges through a classroom experiment

**DOI:** 10.1101/2025.07.12.664517

**Authors:** Huifang Ma, BIOSTAT 824 Student Consortium, Zhicheng Ji

## Abstract

Large language models have shown remarkable capabilities in algorithm design, but their effectiveness in solving data science challenges remains poorly understood. We conducted a classroom experiment in which graduate students used large language models (LLMs) to solve biomedical data science challenges on Kaggle. While their submissions did not top the leaderboards, their prediction scores were often close to those of leading human participants. LLMs frequently recommended gradient boosting methods, which were associated with better performance. Among prompting strategies, self-refinement, where the LLM improves its own initial solution, was the most effective, a result validated using additional LLMs. These findings demonstrate that LLMs can design competitive machine learning solutions, even when used by non-experts.

## Introductions

Large language models (LLMs) are advanced deep learning architectures trained on vast corpora of textual data to understand and generate human-like language. These models, such as GPT^1^, BERT^2^, and their successors, leverage transformer-based architectures to capture complex linguistic patterns, semantic relationships, and contextual dependencies across long sequences of text. LLMs have demonstrated remarkable performance across a wide range of natural language processing tasks. Their generalization ability, combined with scalability through transfer learning and prompt engineering, has positioned LLMs as foundational tools in modern artificial intelligence research and applications.

LLMs have emerged as powerful tools for code generation, transforming natural language instructions into syntactically valid and semantically meaningful code across a variety of programming languages. Early advances were driven by models such as OpenAI’s Codex^3^, which demonstrated the ability to solve around 28.8% of Python problems in the HumanEval benchmark when prompted with a single solution, and up to 70% with multiple samples, outperforming GPT-3 by a significant margin. Building on this foundation, AlphaCode by DeepMind achieved human-level performance in programming contests, placing within the top 54.3% on Codeforces and demonstrating the potential of LLMs to solve complex, competitive tasks^4^. Meta AI’s InCoder^5^ and CodeGen^6^ further advanced LLM capabilities by enabling code infilling and supporting multilingual programming, thereby improving usability for real-world development. SantaCoder^7^ introduced efficient, open-access training for multilingual code generation. StarCoder^8^, trained on over 1 trillion tokens of code, showed strong performance across multiple benchmarks while remaining accessible for academic use. Code Llama, Meta’s most recent release, improved on previous models in both zero-shot and few-shot settings^9^, while WizardCoder^10^ applied instruction tuning to further enhance accuracy. In addition, LLMs like PanGu-Coder2^11^ and DeepSeek-Coder^12^ have demonstrated that instruction tuning, multi-round prompting, and domain-specific adaptation can yield substantial improvements, with some open-source models approaching GPT-4-level performance in controlled benchmarks. Collectively, these findings highlight not only the rapid evolution of LLMs for code generation but also their increasing accessibility, efficiency, and utility in real-world programming tasks. More recent evaluations confirm that state-of-the-art models can match or surpass most human programmers. For example, GPT-4 was found to outperform approximately 85% of participants on LeetCode and GeeksforGeeks coding challenges when optimally prompted^13^.

However, most existing research has focused on algorithm design tasks, such as those found in HumanEval and LeetCode, where LLMs are evaluated based on their ability to produce correct and efficient implementations of well-defined programming problems. In contrast, the performance of LLMs in broader code generation applications remains largely underexplored. One important yet understudied application is their use in solving data science challenges. Unlike algorithm design tasks, which emphasize precise logic, data structures, and deterministic correctness, data science involves building end-to-end analytical pipelines using real-world datasets. These tasks typically require data cleaning, feature engineering, model selection, and performance optimization, often under noisy or imperfect data conditions. Success in data science challenges, such as those hosted on Kaggle, is measured not by exact functional correctness but by the quality of predictions on unseen data, evaluated using metrics like AUC, RMSE, or log loss. This distinction implies that LLMs applied to data science must demonstrate not only programming competence but also the ability to reason about data, select appropriate methods, and adapt code dynamically across diverse contexts. As such, evaluating LLMs in this setting provides a more comprehensive view of their practical utility for real-world problem-solving.

Another limitation of existing research on LLMs concerns the nature of the evaluators themselves. Most benchmarking studies are conducted by researchers who are highly familiar with LLMs, many of whom are experts in prompt engineering or directly involved in LLM development. These individuals tend to craft carefully optimized prompts that leverage deep knowledge of model behavior, often resulting in inflated performance metrics that do not generalize to typical users. In contrast, the broader population of users, such as data scientists, students, or professionals in applied domains, often lack specialized training in how to interact effectively with LLMs. As a result, the actual performance of LLMs when used by non-experts may differ substantially from what published benchmarks suggest. This gap raises concerns about ecological validity and limits the generalizability of current findings to real-world settings.

In this study, we present the results of a classroom experiment in which 33 graduate students enrolled in a data science course were tasked with solving biomedical data science challenges published on Kaggle using OpenAI’s LLMs. We compared the performance of LLM-assisted students with that of human data scientists who participated in the same Kaggle competitions. Additionally, we investigated factors that contribute to the improved performance of LLMs in solving data science problems. Our findings show that the performance of LLMs, when prompted by graduate students, is comparable to that of the majority of human participants. Notably, we found that the use of self-refinement prompting strategies was effective in enhancing model outputs. This study addresses a critical gap by evaluating the ability of LLMs, as used by non-expert users, to solve complex, real-world data science challenges.

## Results

### Experiment design

The classroom experiment was conducted with 33 graduate students enrolled in the course BIOSTAT 824: Case Studies in Biomedical Data Science, offered by the Department of Biostatistics and Bioinformatics at the Duke University School of Medicine during the Spring 2025 semester. As the final project, students were tasked with using OpenAI’s o1 or o3 language model to solve six biomedical data science challenges hosted on Kaggle, a widely used platform for data science competitions. To accommodate the limited computational resources accessible to the students, all six challenges were selected from Kaggle’s “Tabular Playground Series”, where the training datasets are provided in a lightweight, tabular format. To simulate a scenario in which users lack programming expertise, students were restricted to using only natural language prompts when interacting with the LLM. To prevent LLMs from retrieving answers online, students were required to disable the web search function of the OpenAI models. Each student was required to submit the complete chat history and the final code generated for each challenge as part of their project deliverables. The detailed instructions for the final project and the six Kaggle data science challenges are provided in the Methods section.

We then ran the final code submitted by the students to generate the prediction files, submitted these files to the Kaggle website, and recorded the corresponding prediction performance. This performance was compared to that of all human data scientists listed on the leaderboard, and a percentile rank was calculated. Students were graded based on their aggregated performance across all six tasks (Methods).

### Performance overview

Figure 1a shows the evaluation results of all LLM-generated solutions across six Kaggle challenges. In 95% of cases, we successfully generated valid prediction files that passed the Kaggle evaluation system based on the code submitted by the students. In the remaining 5% of cases, the code either contained programming errors or produced invalid prediction files that failed to pass the evaluation. These results suggest that LLMs are highly reliable for real-world applications, as they can autonomously generate functional code that meets external evaluation standards in the vast majority of cases. LLM performance across the six challenges demonstrated a general trend of consistency, with students who performed well on one challenge also tending to perform well on others.

**Figure 1.**
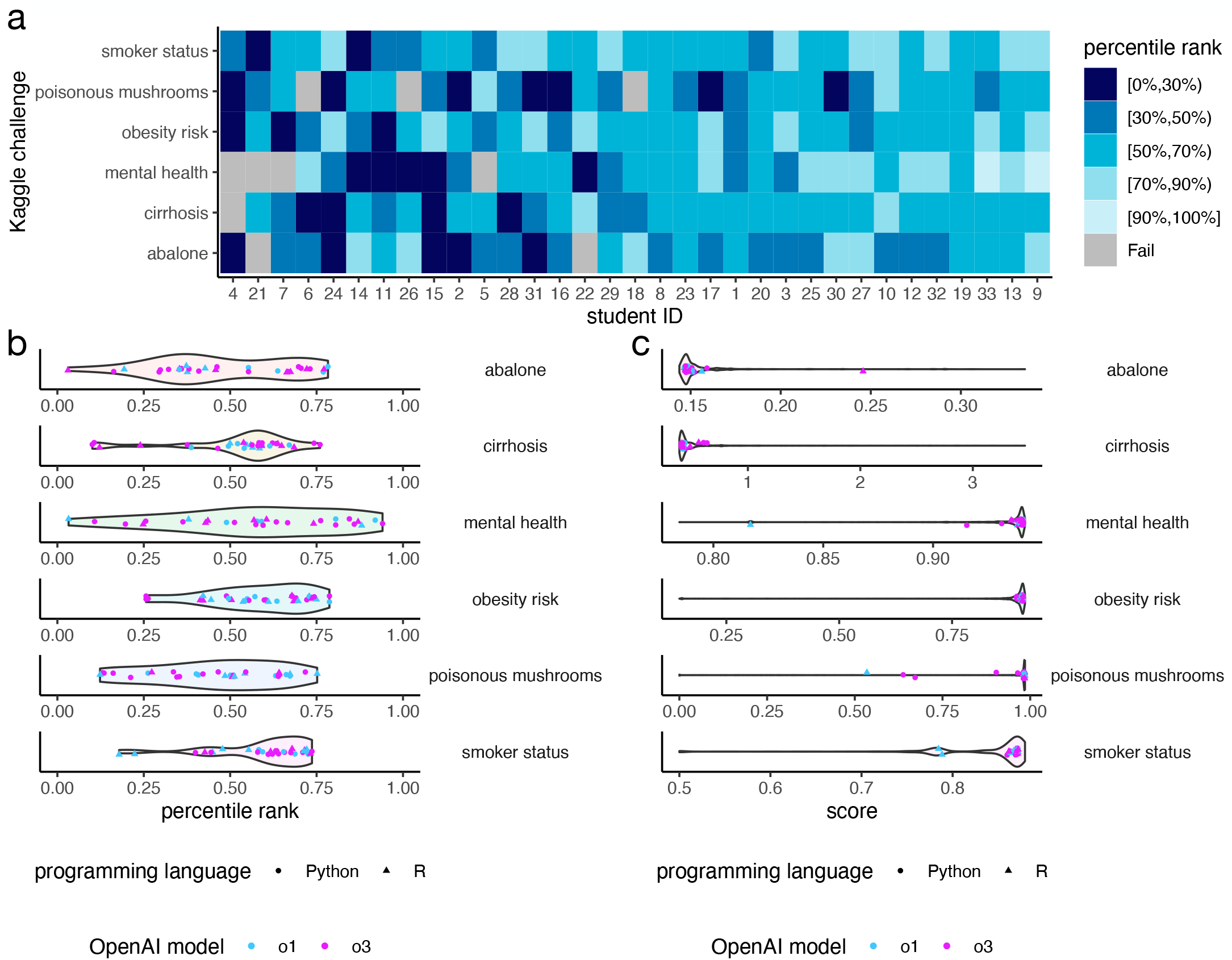
Performance overview. **a**, Kaggle leaderboard percentile rank for each student and each Kaggle challenge. Gray indicates solutions that failed to pass Kaggle’s automatic scoring system. **b**, Violin plots showing the percentile ranks of all student solutions for each Kaggle challenge. Each colored data point represents a single student’s solution. **c**, Violin plots showing the scores of all human participant solutions recorded on the Kaggle leaderboard for each challenge. Each colored data point represents a student’s solution. Note that the violin plots in **b** and **c** represent different populations.

Figure 1b shows the distribution of percentile ranks for the LLM-generated solutions. The majority of solutions achieved a percentile rank between 25% and 75% across all six challenges. There were no substantial differences in performance between solutions generated using different OpenAI models (o1 or o3) or programming languages (R or Python). Figure 1c compares the distribution of performance scores for all human data scientists listed on the Kaggle leaderboard with those of the LLMs. Notably, the absolute performance of the top-performing human data scientists is tightly clustered, meaning that even minor differences in prediction scores can lead to large shifts in percentile rank. As a result, while LLMs’ scores were often very close to those of the best-performing models, their percentile rankings could vary significantly. Nonetheless, most LLM solutions were comparable in accuracy to those of a high-performing subgroup of human data scientists. These results demonstrate that LLMs can generate solutions that are practically effective for a variety of biomedical data science tasks, even if they do not top the leaderboard.

### Machine learning strategies

We manually analyzed the final submission code generated by LLMs and curated the machine learning methods and techniques they employed. Gradient boosting is the most commonly used method, followed by random forest and regression (Figure 2a). On average, gradient boosting appears more than once per submission, as each submission may include multiple software packages implementing different variants, such as XGBoost^14^, CatBoost^15^, and LightGBM^16^. The LLMs’ frequent selection of gradient boosting aligns with its status as a top-performing method in recent machine learning challenges involving tabular and structured data^14^. Neural networks, while widely used in fields such as imaging analysis, are rarely selected by LLMs, likely due to the lightweight and less complex nature of the datasets. Submissions that use gradient boosting achieve significantly better performance than those that do not (Figure 2b), while submissions using random forest or regression tend to perform significantly worse. These findings reinforce the well-established effectiveness of gradient boosting in structured data tasks.

**Figure 2.**
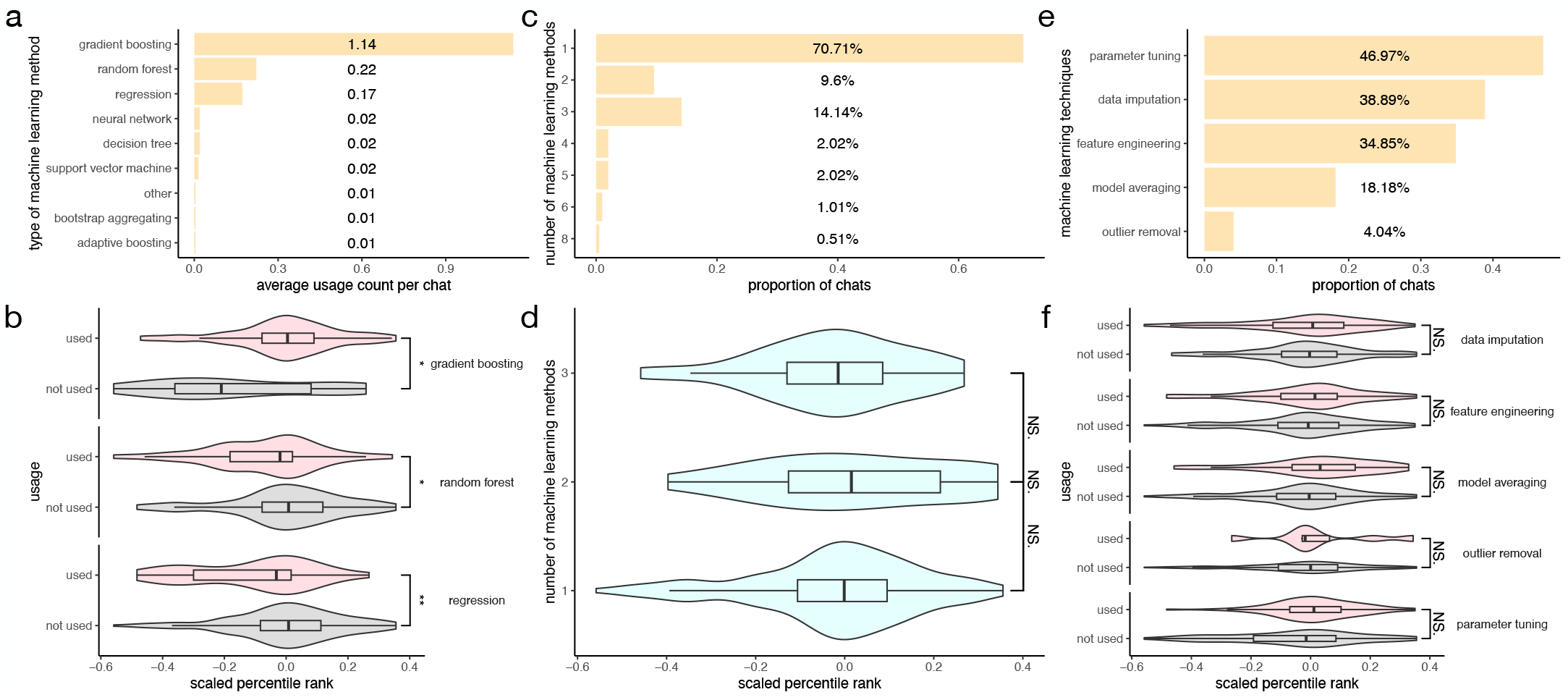
Machine learning strategies. **a**, Average number of uses per chat for different types of machine learning methods. **b**, Distribution of scaled percentile rank comparing solutions that did or did not use a given type of machine learning method. Wilcoxon tests were performed to compare the two distributions. “**” indicates a p-value between 0.001 and 0.01, and “*” indicates a p-value between 0.01 and 0.05. **c**, Proportion of chats that used different numbers of machine learning methods. **d**, Distribution of scaled percentile rank comparing solutions that used different numbers of machine learning methods. Wilcoxon tests were performed to compare the distributions. “NS.” indicates the p-value is not statistically significant. **e**, Proportion of chats that used different machine learning techniques. **f**, Distribution of scaled percentile rank comparing solutions that did or did not use a given machine learning technique. Wilcoxon tests were performed to compare the distributions. “NS.” indicates the p-value is not statistically significant.

Most submissions involve only one type of machine learning method (e.g., gradient boosting), and nearly all include no more than four types (Figure 2c), suggesting that LLMs tend to favor simpler modeling pipelines with a limited set of familiar algorithms rather than constructing complex ensembles. However, performance does not vary substantially with the number of machine learning methods used (Figure 2d), indicating that increasing model diversity alone does not necessarily improve outcomes. Among machine learning techniques, parameter tuning is the most commonly used, appearing in nearly half of the submissions (Figure 2e), reflecting LLMs’ recognition of the importance of hyperparameter optimization. Data imputation, feature engineering, and model averaging are also relatively common, showing awareness of standard preprocessing and ensembling strategies. In contrast, outlier removal is the least frequently used technique, suggesting either a lower emphasis on data cleaning or limitations in the prompt-driven setup regarding quality control. While most techniques are associated with improved performance, the gains are not statistically significant (Figure 2f), implying that the value of these techniques may depend heavily on context and dataset characteristics.

### Prompt strategies

We next manually analyzed the prompt messages and curated the prompt strategies used by the students. Figure 3a shows the proportion of cases in which different prompt strategies were employed. Here, a case refers to a complete chat history from one student addressing one Kaggle challenge. Iterative debugging, which involves repeatedly providing error messages to the LLMs and allowing them to fix errors in the code, is the most commonly used strategy. This suggests that while LLMs are prone to generating programming errors, they also possess a strong ability to correct their own mistakes, as indicated by the low percentage of final answers that failed to pass Kaggle’s evaluation system (Figure 1a). In more than half of the cases, the name of a machine learning method or technique, such as those listed in Figure 2, was mentioned in the chat. Self-refinement, a less commonly reported and studied prompt strategy, also appeared in about half of the chats. Self-refinement refers to prompt messages that ask LLMs to improve their previous response without specifying how the improvement should be made. An example of such a prompt is, “The performance is not good enough, further improve your answer.”

**Figure 3.**
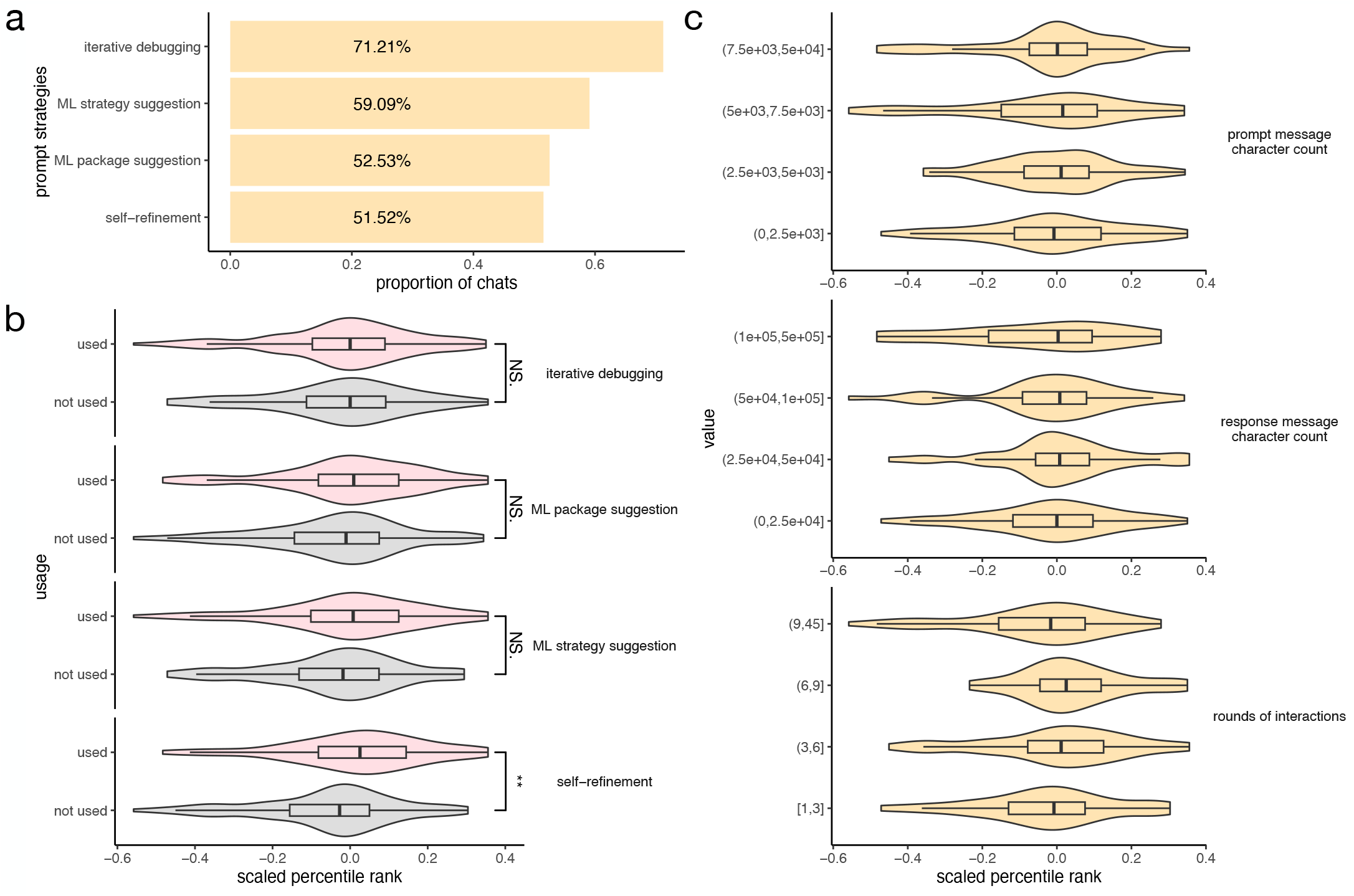
Prompt strategies. **a**, Proportion of chats that used different prompt strategies. **b**, Distribution of scaled percentile rank comparing solutions that did or did not use a given prompt strategy. Wilcoxon tests were performed to compare the distributions. “**” indicates a p. value between 0.001 and 0.01, and “NS.” indicates the p-value is not statistically significant. **c**, Distribution of scaled percentile rank comparing chats across different intervals of character count in the prompt message, response message, and number of interaction rounds.

Figure 3b compares the changes in performance when a specific prompt strategy is used versus not used. Only self-refinement substantially improves performance, while iterative debugging, ML package suggestion, and ML strategy suggestion do not lead to significant improvements. These results highlight the unique contribution of self-refinement to the effectiveness of LLM-assisted code generation. Unlike strategies that provide concrete feedback such as error messages or package suggestions, self-refinement prompts engage the LLMs in a more autonomous optimization process. This may encourage the model to explore a broader solution space or refine its reasoning, leading to more effective model choices or better hyperparameter tuning. The lack of significant improvement from other strategies suggests that simply providing error feedback or naming ML tools does not consistently lead to better outcomes, possibly because such information is already well-handled by the LLM without needing explicit instruction. Overall, these findings emphasize the value of prompting LLMs with open-ended, outcome-oriented instructions that allow them to self-direct their improvements.

We also found that other factors in a chat history, including the total number of characters in the prompt or response messages and the number of interaction rounds, have only a minor influence on performance (Figure 3c). These findings suggest that longer or more interactive conversations with LLMs do not necessarily lead to better performance. This implies that the quality and type of prompt strategies, rather than the quantity of text or rounds of interaction, are more critical in influencing outcomes.

Finally, we evaluated the effectiveness of the self-refinement strategy on three LLMs that were not used in the classroom experiment: GPT-4o, Gemini 2.5 Flash, and Claude Sonnet 4. For each of the six Kaggle data science challenges, each LLM was prompted to generate an initial solution and then produce a self-refined version (Methods). In nearly 90% of the cases, the self-refined solution outperformed the initial one and led to an average improvement of 11.5% in percentile rank (Figure 4). These results align with the classroom experiment findings and suggest that the self-refinement strategy can be broadly applied to improve performance in data science challenges across different LLMs.

**Figure 4.**
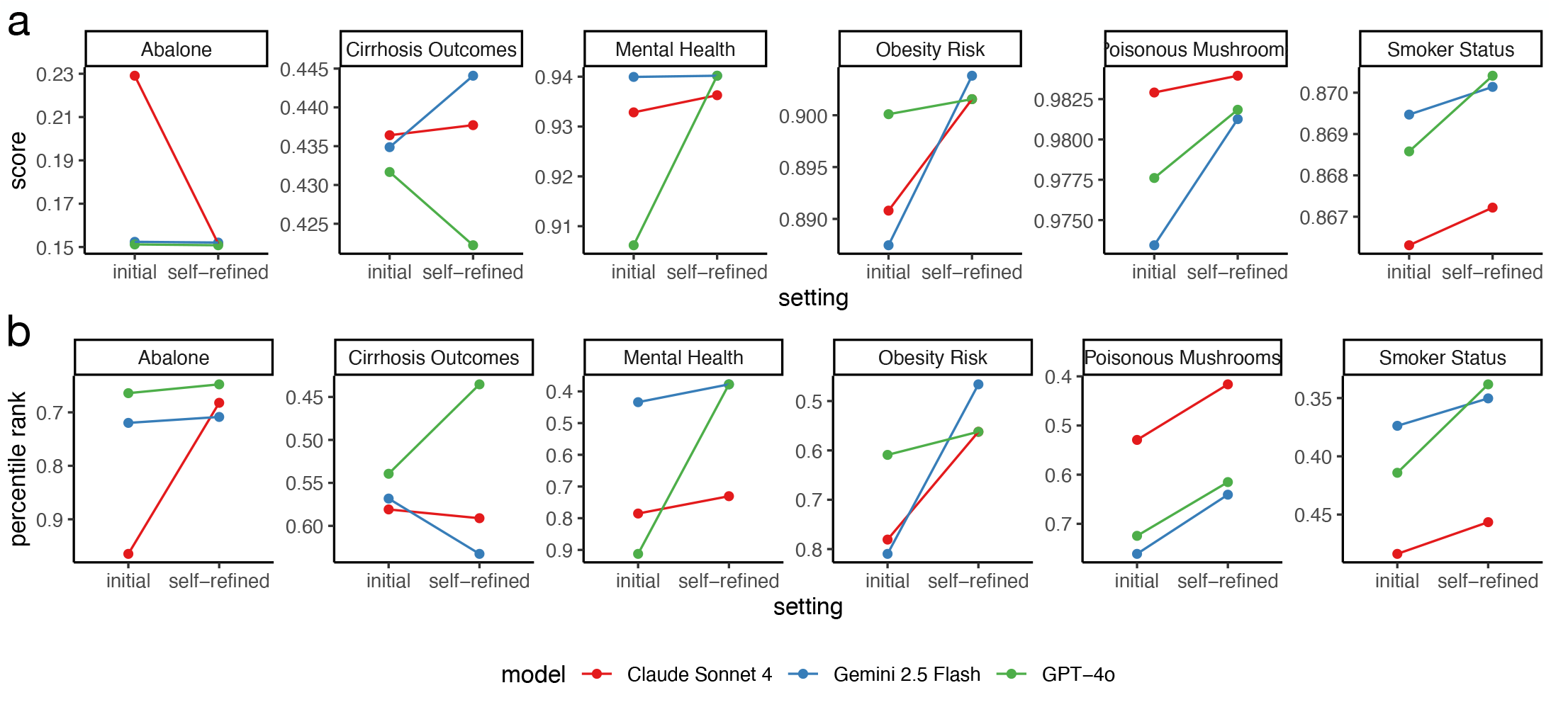
Validation of self-refinement. **a-b**, Score (**a**) and percentile rank (**b**) comparing the initial and self-refined solutions.

## Discussions

We presented results from a classroom experiment in which graduate students used LLMs to participate in biomedical data science challenges hosted on Kaggle. We found that while students’ submissions did not top the leaderboards, their prediction scores were often close to those of top-performing human data scientists. LLMs tended to favor gradient boosting methods, and solutions incorporating these methods generally outperformed those that did not. Among various prompting strategies, the self-refinement approach, where the LLM is prompted to improve upon its own initial solution, proved to be the most effective. The effectiveness of this strategy was further validated using LLMs not included in the classroom setting. Overall, this study demonstrates that LLMs can reliably design machine learning algorithms with performance comparable to that of experienced human data scientists, even when operated by users without specialized expertise in LLMs.

This study has several limitations. First, the graduate students who participated in the classroom experiment shared similar academic backgrounds, making it unclear how users with more diverse training might perform. Second, all Kaggle challenges used in this study involved tabular prediction tasks within the domain of biomedical data science, reflecting the theme of the course and constraints on computational resources. It remains unknown how LLMs would perform on challenges involving larger or more complex datasets. Finally, LLMs were restricted from querying online resources, and students were not allowed to provide programming code to ensure a fair evaluation. How LLMs perform when given full autonomy, including access to online content and user-provided code, remains an open question.

## Methods

### Kaggle data science challenges

The following six Kaggle data science challenges were involved in this study:

1. Multi-Class Prediction of Obesity Risk (https://www.kaggle.com/competitions/playground-series-s4e2/).
2. Binary Prediction of Smoker Status using Bio-Signals (https://www.kaggle.com/competitions/playground-series-s3e24/).
3. Multi-Class Prediction of Cirrhosis Outcomes (https://www.kaggle.com/competitions/playground-series-s3e26/).
4. Binary Prediction of Poisonous Mushrooms (https://www.kaggle.com/competitions/playground-series-s4e8/).
5. Regression with an Abalone Dataset (https://www.kaggle.com/competitions/playground-series-s4e4).
6. Exploring Mental Health Data (https://www.kaggle.com/competitions/playground-series-s4e11/).

### Initial instructions for the final project

The following instructions for the final project were given to the students at the beginning of the course:

“You are required to participate in six Kaggle challenges. In the final project, you are NOT allowed to write your own code.

Instead, you are required to provide prompts to OpenAI’s o1 model, which will generate the actual code.

For each task, you need to submit:

1. Your chat history URL with o1.
2. The code generated by o1. This should be the exact output from the o1 model and must be included in the chat history (which will be validated). This code will be used to evaluate performance. It is not required to be the last code generated by o1 in the entire chat history. It can be code generated earlier in the conversation. For instance, o1 may have generated some other code later, but you decided the previous code was better. The input can only be “train.csv” and “test.csv” provided by Kaggle.
3. A brief description of your prompt strategy and what you think contributed to the improvement of o1’s performance (you can provide one document for all six challenges).

When communicating with o1, you are only allowed to use natural language to instruct o1. Images are not permitted as input. You may recommend the name of the model, software package, or provide other useful information to o1, but only in natural language. Programming language is not restricted. You are prohibited from providing any actual programming code, or natural language directly translated from programming code (e.g., pseudocode) to o1. You will be notified and given a chance to fix it within a short timeframe if this happens. Violations will result in a score of 0 for the specific challenge and will disqualify you from the in-class ranking for the specific challenge and potential authorship of the paper.

For each challenge, you will receive a score of:

- 0 if any of the three required documents are missing, if the rules are violated, or if there is a bug in the final code that prevents obtaining a valid Kaggle ranking.
- 3.0 if you submit all three documents and the code is valid, resulting in a valid Kaggle ranking (regardless of percentile).
- 3.3 if you submit all three documents and your solution performs better than 30% of human participants on Kaggle.
- 3.5 if you submit all three documents and your solution performs better than 50% of human participants on Kaggle.
- 3.7 if you submit all three documents and your solution performs better than 70% of human participants on Kaggle.
- 4.0 if you submit all three documents and your solution performs better than 90% of human participants on Kaggle.

Bonus for each challenge:

- +1.0 if you rank first in class.
- +0.6 if you rank 2-3 in class.
- +0.3 if you rank 4-5 in class.

Each of the six challenges will be graded separately. The final score will be an average of the six scores for the six Kaggle challenges.”

### Updated instructions for the final project

The following instructions for the final project were given to the students after OpenAI replaced o1 with the o3 model:

“OpenAI has updated its model list, introducing the new o3 model while removing the older o1 model. Consequently, the rules for the final project have been revised.

You may complete the project with either the o1 or o3 model. Results from both models will be evaluated together, with no score adjustment to favor one over the other, so choosing o1 carries the risk that o3 may outperform it. If you opt for the o3 model, do not use the “Search the web”, “Deep research”, or “Canvas” functions. Usage of these features can be verified in your chat history. Projects that employ them will require resubmission.

All other project guidelines remain unchanged.

For those of you using o3, the model may still perform a web search even when the “Search the web” button is not toggled. In that case, disable web search in the settings: Click your user icon in the top right corner of the page, select “Customize ChatGPT”, scroll down to “Advanced”, and uncheck “Web search”. This will disable web search at the system level. Note: changing the user settings is not required. If your response does not involve web browsing in the first place, please ignore this and you don’t need to redo anything.

Please double check that web browsing is not used by the o3 or o1 model in your response, as doing so violates the project rules. To verify, click “Thought for XX seconds” to view the step by step reasoning, and make sure no step shows “Searched the web” or something similar. If o3 or o1 performs web browsing in your submission, we will return it and ask you to redo it.”

### Evaluation of self-refinement strategy

We prompted the online versions of GPT-4o, Gemini 2.5 Flash, and Claude Sonnet 4 to generate programming code for solving the same six Kaggle data science challenges used in the classroom experiment. For the initial query, the prompt message consisted of the following components:

- “Write code for performing the following machine learning challenge:”
- Information about the challenge copied from the Kaggle website.
- The first 10 lines of the training file, test file, and example submission file.
- “Generate code in a single chunk. Do not provide explanations. The code cannot take longer than 1 hour to finish.”

For the self-refinement query, the prompt message is as follows: “The performance is not good enough. Further improve your answer.”

### Calculation of scaled percentile rank

We introduced the scaled percentile rank to account for differences in percentile rank distributions across Kaggle challenges, enabling pooled analysis across challenges. For each Kaggle data science challenge, the scaled percentile rank was calculated by subtracting the median percentile rank of all students in that challenge from each student’s raw percentile rank.

## Author contributions

All students in the BIOSTAT 824 Student Consortium participated in the classroom experiment. Z.J. conceived the study. H.M. and Z.J. analyzed the data. Z.J. wrote the manuscript.

BIOSTAT 824 Student Consortium: Tara Al-Hashimy,Austin Allen,Nan Cen,Orlando Chen,Yongyin Chen,Yutian Chen,Tong Cheng,Yueqi Gu,Beijie Ji,Xiaohui Jiang,Fengnan Li,Peiyu Li,Yueshan Liang,Bena Liu,Coco Liu,Elisa Ma,Zhicheng Ma,Vicky Shao,Mengyao Shi,Jiang Shu,Leyi Sun,Rushi Tang,Hanyu Wang,Vivian Wang,Yuxin Wang,Krissie Wilson,Ruobing Xue,Tianyi Yang,Alison Yu,Allison Yuan,Haiqi Zhang,Vera Zhang,Yinuo Zhang.

## Ethics Statement

This study was reviewed by the Duke Health Institutional Review Board (DUHS IRB) and determined not to involve human subjects (Protocol Pro00117372).

